# GSimp: A Gibbs sampler based left-censored missing value imputation approach for metabolomics studies

**DOI:** 10.1101/177410

**Authors:** Runmin Wei, Jingye Wang, Erik Jia, Tianlu Chen, Yan Ni, Wei Jia

## Abstract

Left-censored missing values commonly exist in targeted metabolomics datasets and can be considered as missing not at random (MNAR). Improper data processing procedures for missing values will cause adverse impacts on subsequent statistical analyses. However, few imputation methods have been developed and applied to the situation of MNAR in the field of metabolomics. Thus, a practical left-censored missing value imputation method is urgently needed. We have developed an iterative Gibbs sampler based left-censored missing value imputation approach (GSimp). We compared GSimp with other three imputation methods on two real-world targeted metabolomics datasets and one simulation dataset using our imputation evaluation pipeline. The results show that GSimp outperforms other imputation methods in terms of imputation accuracy, observation distribution, univariate and multivariate analyses, and statistical sensitivity. The R code for GSimp, evaluation pipeline, vignette, real-world and simulated targeted metabolomics datasets are available at: https://github.com/WandeRum/GSimp.

**Author summary:** Missing values caused by the limit of detection/quantification (LOD/LOQ) were widely observed in mass spectrometry (MS)-based targeted metabolomics studies and could be recognized as missing not at random (MNAR). MNAR leads to biased parameter estimations and jeopardizes following statistical analyses in different aspects, such as distorting sample distribution, impairing statistical power, etc. Although a wide range of missing value imputation methods was developed for –omics studies, a limited number of methods was designed appropriately for the situation of MNAR currently. To alleviate problems caused by MNAR and facilitate targeted metabolomics studies, we developed a Gibbs sampler based missing value imputation approach, called GSimp, which is public-accessible on GitHub. And we compared our method with existing approaches using an imputation evaluation pipeline on real-world and simulated metabolomics datasets to demonstrate the superiority of our method from different perspectives.

## Introduction

Missing values are commonly observed in mass spectrometry (MS) based metabolomics datasets. Many statistical methods require a complete dataset, which makes missing data an inevitable problem for subsequent data analysis. Generally, there are three types of missing values, missing not at random (MNAR), missing at random (MAR) and missing completely at random (MCAR) [1,2]. Unexpected missing values are considered as MCAR if they originate from random errors and stochastic fluctuations during the data acquisition process (e.g., incomplete derivatization or ionization). MAR assumes the probability of a variable being missing depends on other observed variables [1,2]. Thus, missing values due to suboptimal data preprocessing, e.g., inaccurate peak detection and deconvolution of co-eluting compounds can be defined as MAR. Targeted metabolomics studies have been widely used for the accurate quantification of specific groups of metabolites. Due to the limit of compound quantifications (LOQ), missing values are usually caused by signal intensities lower than LOQ, also known as left-censored missing, which can be assigned to MNAR.

The processing of missing values has been developed and studied in MS data, which is an indispensable step in the metabolomics data processing pipeline [3]. One simple but naïve solution is the substitution of missing by determined values, such as zero, half of the minimum value (HM) or LOQ/c where c denotes a positive integer. Determined value substitutions, although commonly applied for dealing with missing values in metabolomics studies [4–6], can significantly affect the subsequent statistical analyses in different ways, e.g. underestimate variances of missing variables, decrease statistical power, fabricate pseudo-clusters among observations, etc. [1]. Advanced statistical imputation methods have been developed for –omics studies, e.g., k-nearest neighbors (kNN) imputation [7], singular value decomposition (SVD) imputation [8,9], random forest (RF) imputation [10]. Several metabolomics data analysis software tools provide different methods of dealing with missing values [11–15]. MetaboAnalyst [15–17], one widely used metabolomics analysis toolkit, provides Probabilistic PCA (PPCA), Bayesian PCA (BPCA) and SVD imputation. However, these methods are mainly aiming at imputing MCAR/MAR and not suitable for the situation of MNAR. A limited number of approaches dealing with left-censored missing values were applied by researchers [18,19]. Quantile regression approach for left-censored missing (QRILC) imputes missing data using random draws from a truncated distribution with parameters estimated using quantile regression [20]. Although this imputation keeps the overall distribution of missing parts compared to determined value substitutions, it may produce random results since no more information is used for the prediction of missing parts. Another imputation method recently developed for MNAR is k-nearest neighbor truncation (kNN-TN) by Shah, et al. [21]. This approach applies Maximum Likelihood Estimators (MLE) for the means and standard deviations of missing variables based on truncated normal distribution. Then a Pearson correlation based kNN imputation method was implemented on standardized data. Although the author stated that kNN-TN could impute both MNAR and MAR, the imputed values were entirely dependent on the nearest neighbors while no constraint was placed upon the imputation. Thus, this approach might cause an overestimation of missing values.

To reduce adverse effects caused by missing values during metabolomics data analyses, we developed a left-censored missing value imputation framework, GSimp, where a prediction model was embedded in an iterative Gibbs sampler. We then compared GSimp with HM, QRILC, and kNN-TN on two real-world metabolomics datasets and one simulation dataset to demonstrate the advantages of GSimp regarding imputation accuracy, observation distribution, univariate analysis, multivariate analysis and sensitivity. Our findings indicate that GSimp is a robust method to handle left-censored missing values in targeted metabolomics studies.

## Results

### Gibbs sampler in GSimp

A variable containing missing elements from FFA dataset was randomly selected to track the sequence of corresponding parameters and estimates across the first 500 iterations out of a total of 2000 (100 × 20) iterations using GSimp. From Fig 1, we can observe that both fitted value 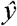 and sample value 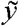 reach to the convergence after iterations and the standard deviation estimate σ drop to a steady state with small values. In addition, an upper constraint for the distribution of 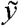 indicated that it was drawn from a truncated normal distribution.

**Fig 1.**
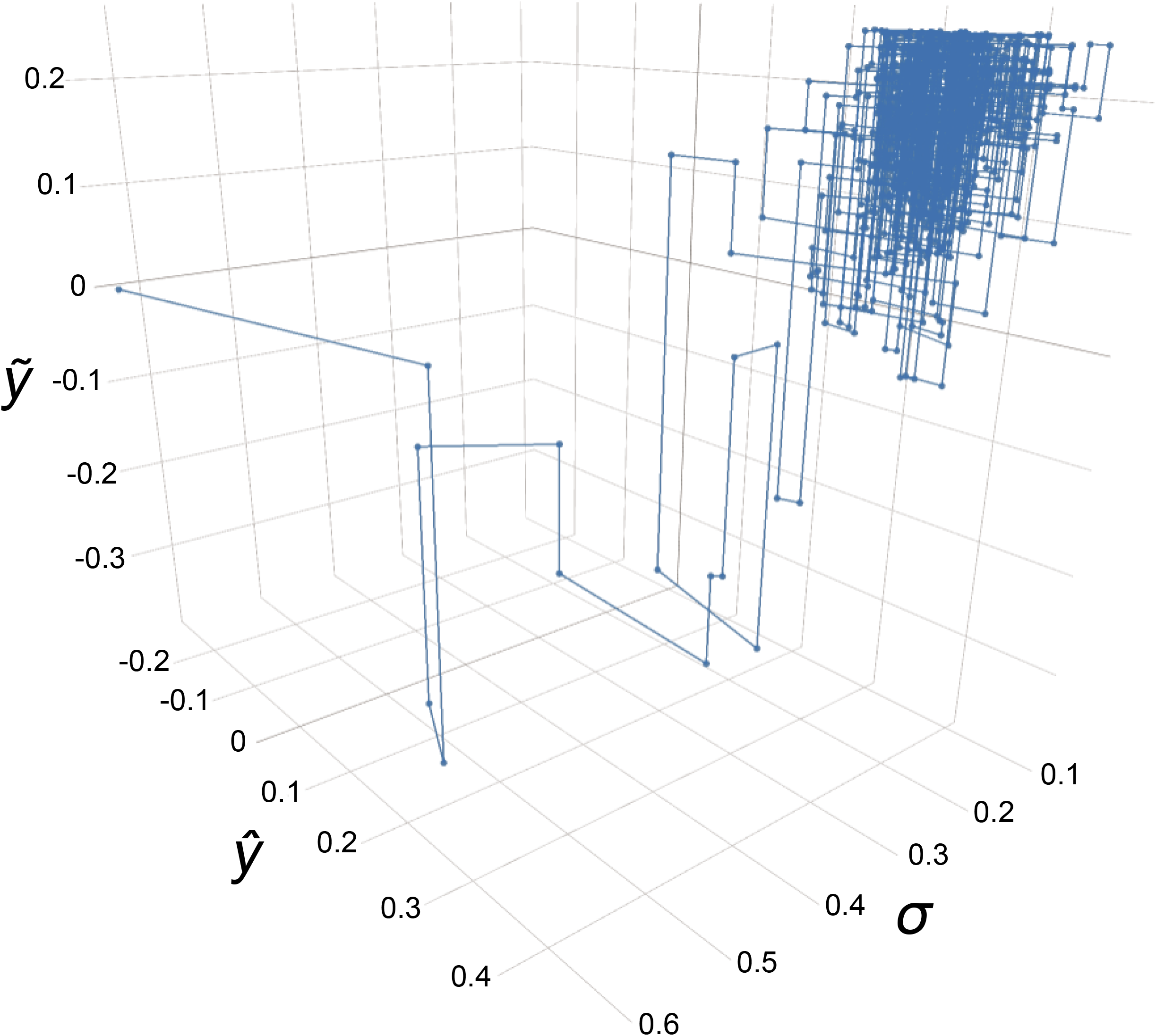
Sequentially parameters updating in GSimp. The first 500 iterations out of a total of 2000 (100×20) iterations using GSimp where 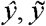 and σ represent fitted value, sample value and standard deviation correspondingly.

### Imputation comparisons

We evaluated four different MNAR imputation/substitution methods on FFA, BA targeted metabolomics and simulation datasets. First, we measured the imputation performances using label-free approaches. SOR was used to measure the imputation accuracy regarding the imputed values of each missing variable. From the upper panel of Fig 2, we can observe that GSimp has the best performance with the lowest SOR across all varying numbers of missing variables in both FFA and BA datasets. To measure the extent of imputation induced distortion on observation distributions, the PCA-Procrustes analysis was conducted between the original data and imputed data. The lower panel of Fig 2 shows that GSimp has the lowest Procrustes sum of squared errors compared to other methods, which means GSimp kept the overall observation distribution of original dataset with the least distortions.

**Fig 2.**
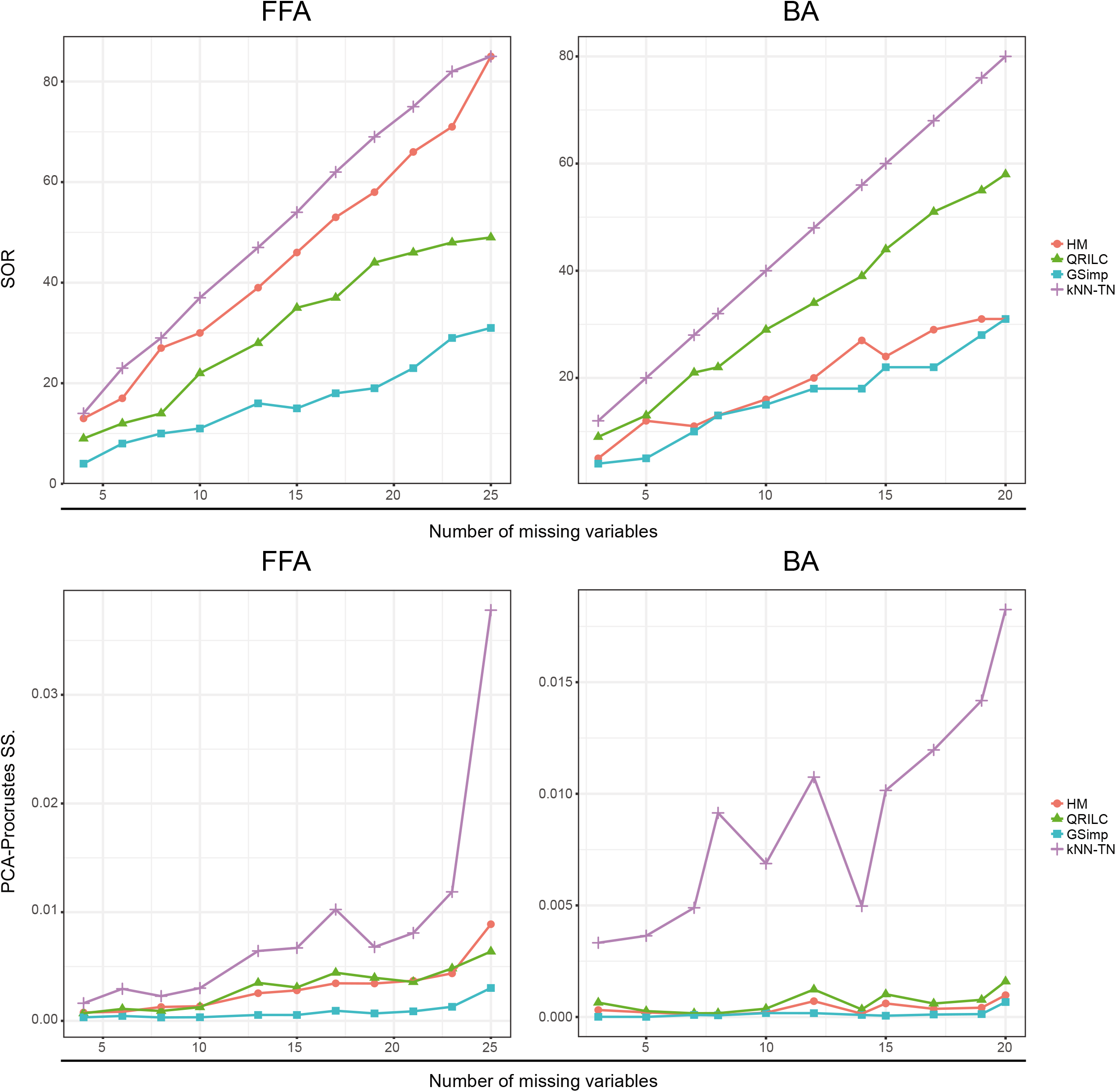
Evaluations of different imputation methods using unlabeled approaches. SOR on FFA dataset (upper left) and BA dataset (upper right) along with different numbers of missing variables based on four imputation methods: HM (red circle), QRILC (green triangle), GSimp (blue square), and kNN-TN (purple cross). PCA-Procrustes sum of squared errors on FFA dataset (lower left) and BA dataset (lower right) along with different numbers of missing variables based on four imputation methods: HM (red circle), QRILC (green triangle), GSimp (blue square), and kNN-TN (purple cross).

Then, we measured the imputation performances with binary labels provided. We compared the results of univariate and multivariate analyses for imputed and original datasets. Since this is a case-control study, student’s *t*-tests were applied for univariate analyses. Then we compared the results by calculating Pearson’s correlation between log-transformed *p*-values calculated from imputed and original data for missing variables. Again, GSimp performs best with the highest correlations among four methods (upper panel of Fig 3) along with different numbers of missing variables, and it implies GSimp keeps the most biological variations regarding the univariate analyses results. For the multivariate analyses, we applied PLS-DA to distinguish the group differences. Similarly, we conducted PLS-Procrustes analysis while PLS was employed as a supervised dimension reduction technique. The lower panel of Fig 3 demonstrates that GSimp preferably restores the original observation distribution with the lowest Procrustes sum of squared errors among four imputation methods.

**Fig 3.**
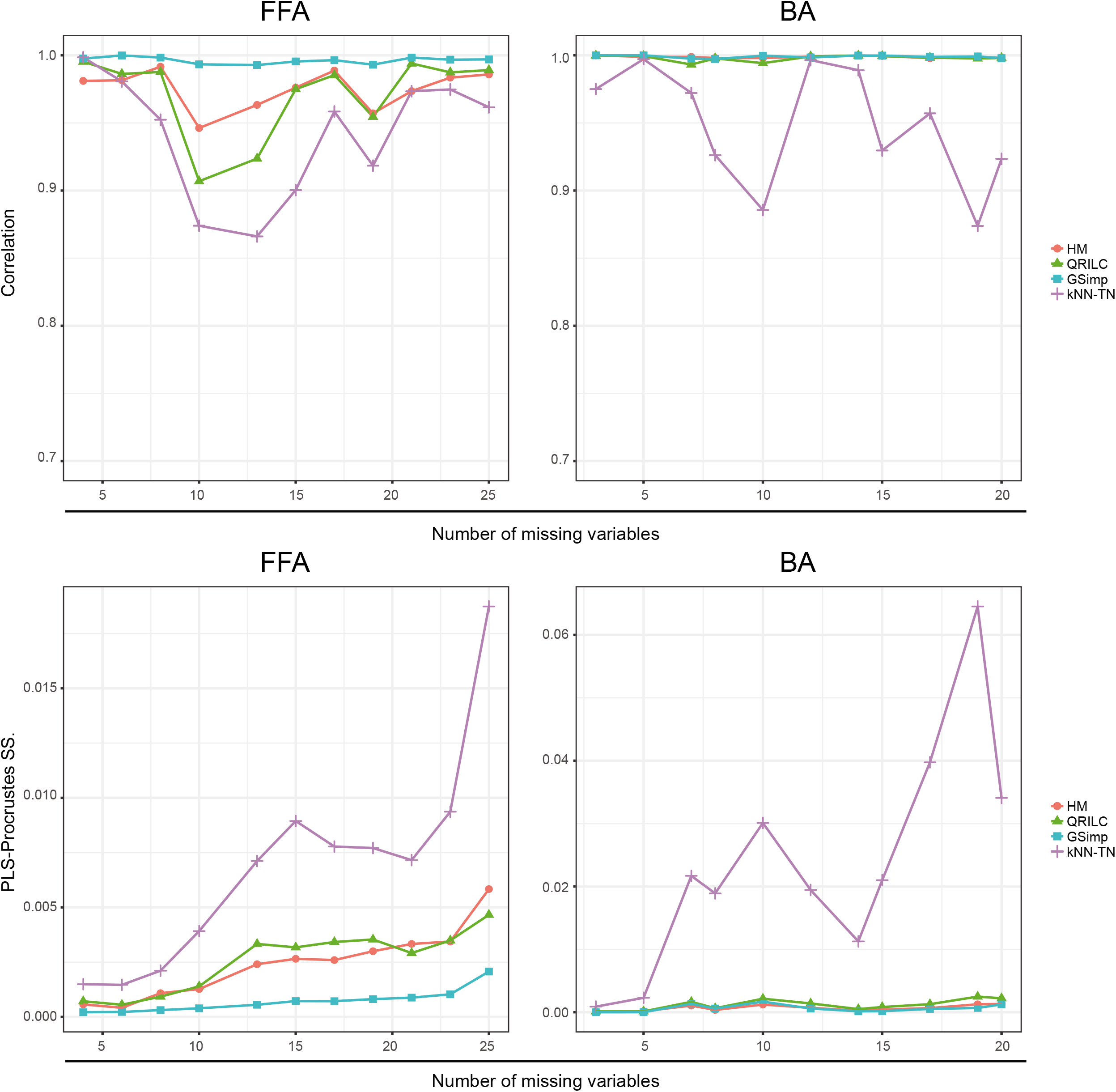
Evaluations of different imputation methods using labeled approaches. Pearson’s correlation between log-transformed p-values of student’s t-tests on FFA dataset (upper left) and BA dataset (upper right) along with different numbers of missing variables based on four imputation methods: HM (red circle), QRILC (green triangle), GSimp (blue square), and kNN-TN (purple cross). PLS-Procrustes sum of squared errors on FFA dataset (lower left) and BA dataset (lower right) along with different numbers of missing variables based on four imputation methods: HM (red circle), QRILC (green triangle), GSimp (blue square), and kNN-TN (purple cross).

On the simulation dataset, we compared QRILC, kNN-TN, and GSimp using same approaches. Consistent results were recognized (S1 Fig), and GSimp presents the best performances on the simulation dataset with the lowest SOR and PCA/PLS-Procrustes sum of squared errors and the highest correlation of univariate analysis results. Moreover, to examine the influences of statistical power using different imputation methods, we calculated *TPR* as the capacities to detect differential variables on different imputation datasets. Again, with both *p*-cutoff of 0.05 and 0.01, GSimp shows the overall highest *TPR* over different missing numbers (Fig 4). This implies that GSimp impairs the sensitivity to the least extent among three methods, which is reasonable since GSimp also keeps the highest correlation of *p*-values in previous comparisons.

**Fig 4.**
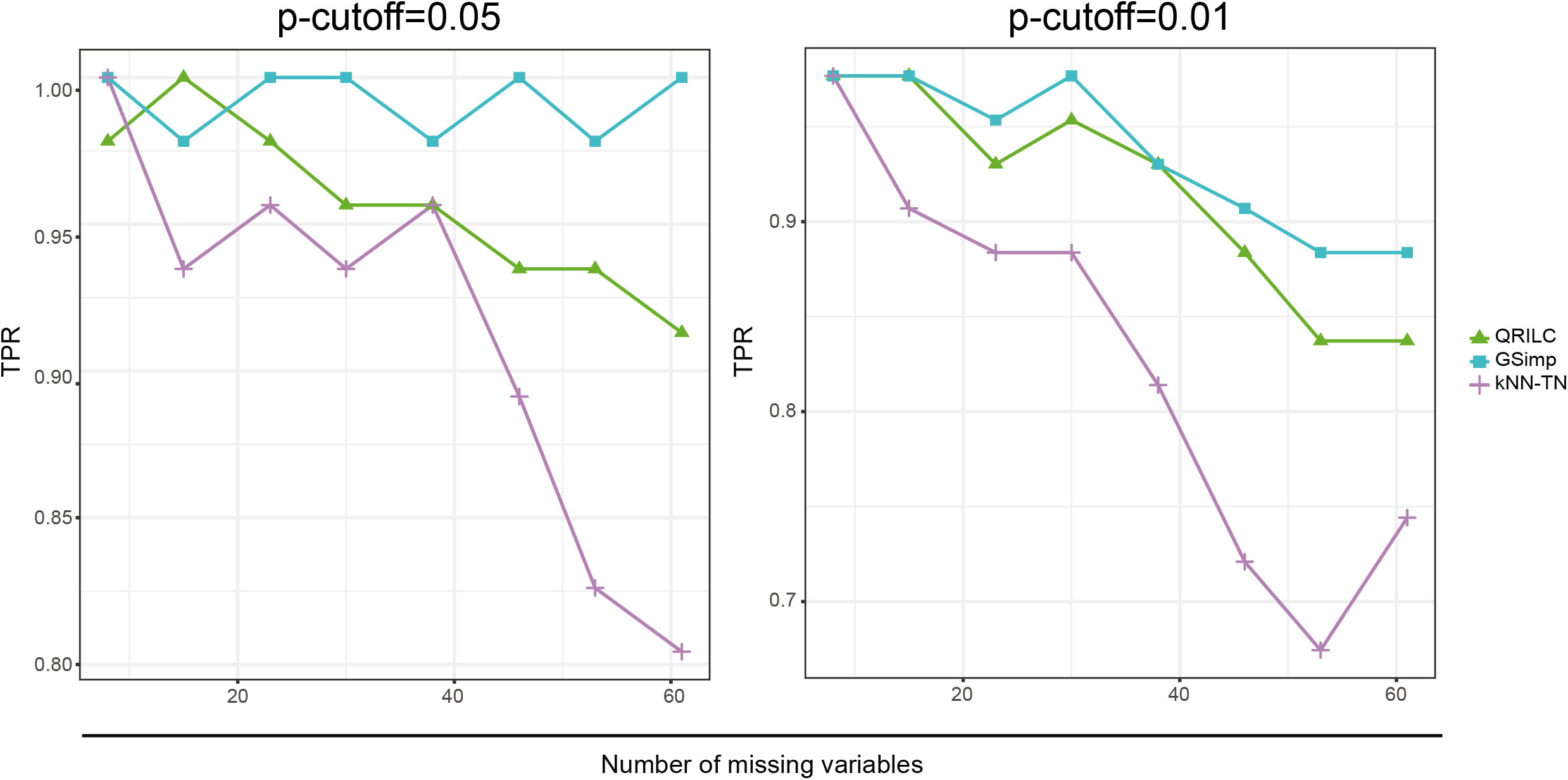
Evaluations of different imputation methods using TPR for various p-cutoffs on simulation dataset. TPR along with different numbers of missing variables based on three imputation methods: QRILC (green triangle), GSimp (blue square), and kNN-TN (purple cross) among different p-cutoff=0.05 (left panel), and 0.01 (right panel).

## Discussion

The purpose of this study is to develop a left-censored missing value imputation approach for targeted metabolomics data analysis. We evaluated GSimp with other three imputation methods (a.k.a kNN-TN, QRILC, and HM) and suggested that GSimp was superior to others using different evaluation methods. To illustrate the performance of GSimp, we randomly selected one variable containing missing values from FFA dataset (Fig 5) to compare the imputed values and original values. Although determined value substitution (e.g. HM) were widely used by researchers in the field of metabolomics, our results indicated that HM could severely distort the data distribution (upper left panel of Fig 5), thus impairing subsequent analyses. In comparison, QRILC kept the overall data distribution and variances (upper right panel of Fig 5). However, random values could be generated by this approach since QRILC imputes each missing variable independently without utilizing the predictive information from other variables. Statistical learning based method, kNN-TN, applied a correlation based kNN algorithm with parameters of missing variables estimated with truncated normal distributions. This method utilized the information of highly correlated variables of targeted missing variable, thus kept a linear trend between original values and imputed values. However, since no constraint was applied for the imputation, a right shift of missing part might occur, causing imputed values to exceed the truncation point (lower left panel of Fig 5). In contrast, GSimp utilized the predictive information of other variables by employing a prediction model and held a truncated normal distribution for each missing element simultaneously, which ensured a favorable linear trend between imputed and original values as well as a reasonable bound for the imputed values (lower right panel of Fig 5).

**Fig 5.**
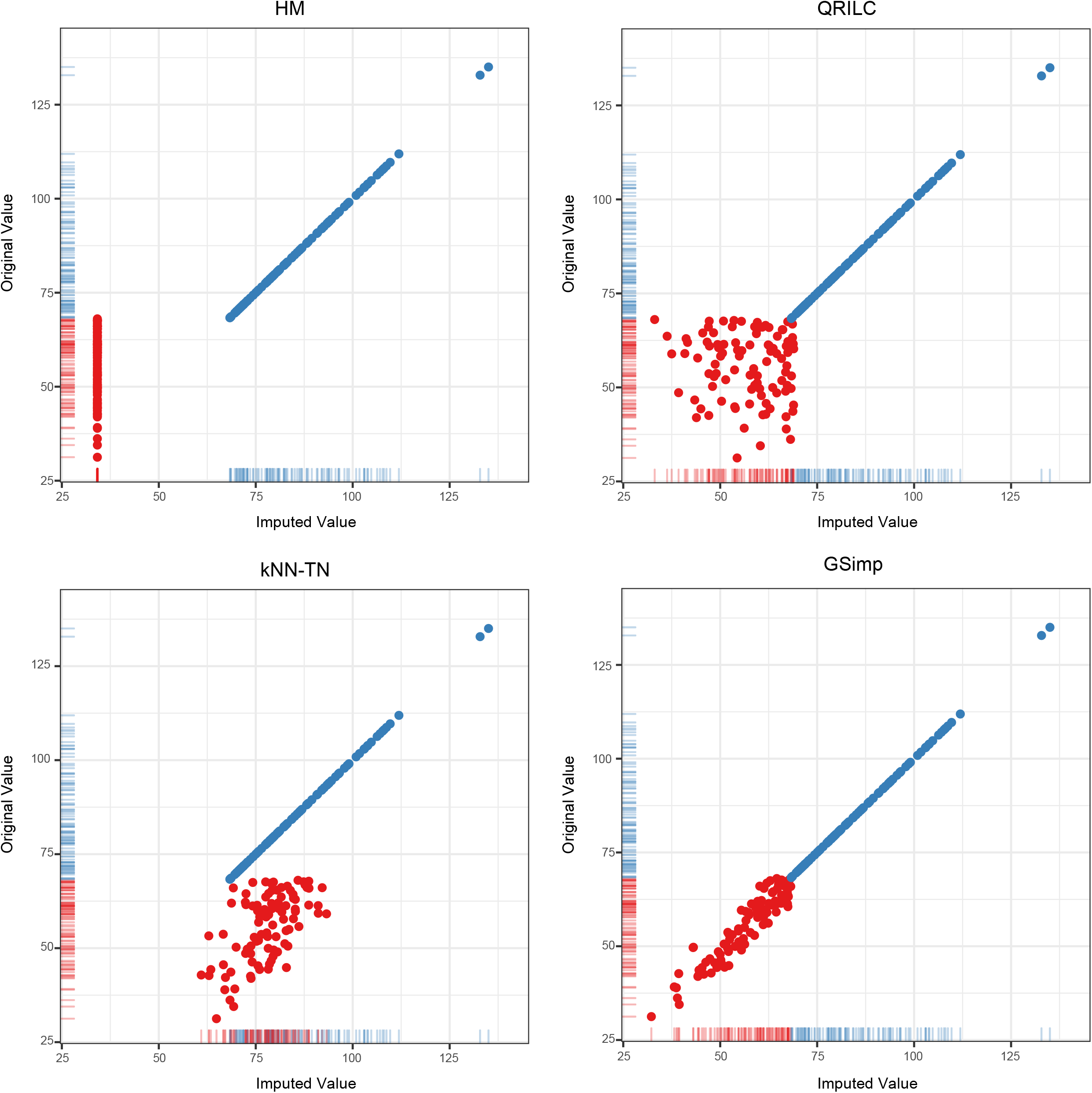
Comparisons of imputed values and original values on an example variable. Scatter plots of imputed values (X-axis) and original values (Y-axis) on one example missing variable while non-missing elements represented as blue dots and missing elements as red dots based on four imputation methods: HM (upper left), QRILC (upper right), kNN-TN (lower left), and GSimp (lower right). Rug plots show the distributions of imputed values and original values.

In our approach, truncated normal distribution was used for the constraint of imputation results in Gibbs sampler steps. We applied the minimum observed value of missing variable as an informative upper truncation point and -∞ as a non-informative lower truncation point considering the situation of left-censored missing. Other values could also be applied in real-world metabolomics analyses, such as a known LOQ of a metabolite can be set as an upper truncation point. Additionally, when signal intensity of certain compound is larger than the upper limit of quantification range or saturation during instrument analysis, an informative lower truncation point could be correspondingly applied for the right-censored missing value. What’s more, when non-informative bounds for both upper and lower limits (e.g., +∞, -∞) were applied, our GSimp could be extended to the situation of MCAR/MAR. With the flexible usage of upper and lower limits, our approach may provide a versatile and powerful imputation technique for different missing types. For other –omics datasets with missing values (especially MNAR), e.g. single cell RNA-sequencing data, we could also apply this method with few modifications of our default settings. Thus, it is worthy to evaluate our approach, GSimp, in other complex scenarios in the future.

Since GSimp employed an iterative Gibbs sampler method, a large number of iterations (*iters_all*=20, *iters_each*=100) are preferable for the convergence of parameters. However, as we tested on the simulation dataset with different number of iterations, a much less iterations (*iters_all*=10, *iters_each*=50) won’t severely affect the imputation accuracy (S2 Fig). Among iterations for the whole data matrix, we applied a sequential imputation procedure for missing variables from the least number of missing values to the most. Such sequential approach improves imputation performances compared to parallel imputation approach.

## Materials and Methods

### Diabetes datasets

We employed datasets from a study of comparing serum metabolites between obese subjects with diabetes mellitus (N=70) and healthy controls (N=130) where N represents the number of observations. Dataset 1: a total of 42 free fatty acids (FFAs) were identified and quantified in those participants in order to evaluate their FFA profiles [22]. Dataset 2: a total of 34 bile acids (BAs) were identified and quantified in a similar way using different analytical protocol [23].

### Simulation dataset

For the simulation dataset, we first calculated the covariance matrix *Cov* based on the whole diabetes dataset (P=76) where P represents the number of variables. Then we generated two separated data matrices with the same number of 80 observations from multivariate normal distributions, representing two different biological groups. For each data matrix, the sample mean of each variable was drawn from a normal distribution *N*(0, 0.5^2^) and *Cov* was kept using SVD. Then, two data matrices were horizontally (column-wise) stacked together as a complete data matrix (N×P=160×76) so that group differences were simulated and covariance was kept.

### MNAR generation

For two real-world targeted metabolomics datasets, we generated a series of MNAR datasets by using the missing proportion (number of missing variables/number of total variables) from 0.1 to 0.6 in a step of 0.05 with MNAR cut-off for each missing variable drawn from a uniform distribution *U*(0.1, 0.5) The elements lower than the corresponding cut-off were removed and replaced with NA. For the simulation dataset, we generated a series of MNAR datasets by using the missing proportion from 0.1 to 0.8 step by 0.1 with MNAR cut-off drawn from *U*(0.3, 0.6) for a more rigorous testing.

### Prediction model

A prediction model was employed for the prediction of missing values by setting a targeted missing variable as outcome and other variables as predictors. Different prediction models, e.g., linear regression, elastic net [24], regression trees [25] and random forest [26], etc. could be embedded in our imputation framework. Elastic net was applied in our approach as an ideal prediction model considering its stability, accuracy, and efficiency. This model is a regularized regression with the combination of L1 and L2 penalties of the LASSO [27] and ridge [28] methods. The estimates of regression coefficients in elastic net are defined as

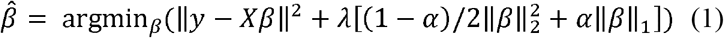

The L2 penalty 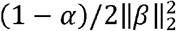 improves the model’s robustness by controlling the multicollinearities among variables which are widely existed in high-dimensional–omics data. And the L1 penalty 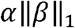 controls the number of predictors by assigning zero coefficients to the “unnecessary” predictors. From a Bayesian point of view, the regularization is a mixture of Gaussian and Laplacian prior distributions of coefficients which can pull the full model of maximum likelihood estimates 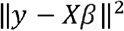 towards the null model of prior coefficients distribution, thus controls the risk of overfitting and increase the model robustness. R package *glmnet* was used for the elastic net. We set hyperparameters λ as 0.01 (default setting for high-dimensional data) and α as 0.5 (an equally mixture of LASSO and ridge penalties) [29].

### Gibbs sampler

Gibbs sampler is a Markov Chain Monte Carlo (MCMC) technique that sequentially updates parameters while others are fixed. It can be used to generate posterior samples. For each missing variable in the dataset, we applied a Gibbs sampler to impute the missing values by sampling from a truncated normal distribution with prediction model fitted value as mean and root mean square deviation (RMSD) of missing part as standard deviation while truncated by specified cut-points. Assuming we have a *n* × *p* data matrix ***X*** = (***X***_1_, ***X***_2_, ***X***_3_, …, ***X****_p_*) with only one variable ***X****_j_* containing left-censored missing values. We denote ***X****_j_* as ***y*** and the missing part as ***y****m* with length *m* and non-missing part as ***y****_f_* with length *f*, and the rest of matrix ***X****_-j_* as ***X****’*. We can then set the lower truncation point *lo* as -∞ (centralized data) or 0 (original data) and upper *hi* as the minimum value of ***y****_f_* or a given LOQ. The truncation bounds ensure imputation results are constrained within [*lo*, *hi*]. Then, the Gibbs sampler approach can be described as following steps:

Step-1 (initialization): we initialize missing values (QRILC in our case), and get ***y****’*;

Step-2 (prediction): we then build a prediction model (elastic net in our case): ***y****’* ~ ***X****’*;

Step-3 (estimation): based on the prediction model, we get the predicted value 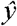 and the root mean square deviation (RMSD) of missing part 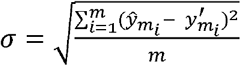 where 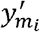 and 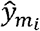 are *i*th initialized/imputed value and fitted value respectively;

Step-4 (sampling): we draw sample 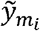 from a truncated normal distribution 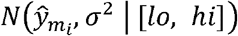 for *i*th missing element and update ***y****’*.

We iteratively repeat step-2 to step-4 and update ***X****_j_*.

### GSimp framework

A whole data matrix ***X*** = (***X***_1_, ***X***_2_, ***X***_3_, …, ***X****_p_*) contains a number of *k* (*k* ≤ *p*) left-censored missing variables. We present our imputation framework as following algorithm.

#### Algorithm Gibbs sampler based left-censored missing value imputation approach

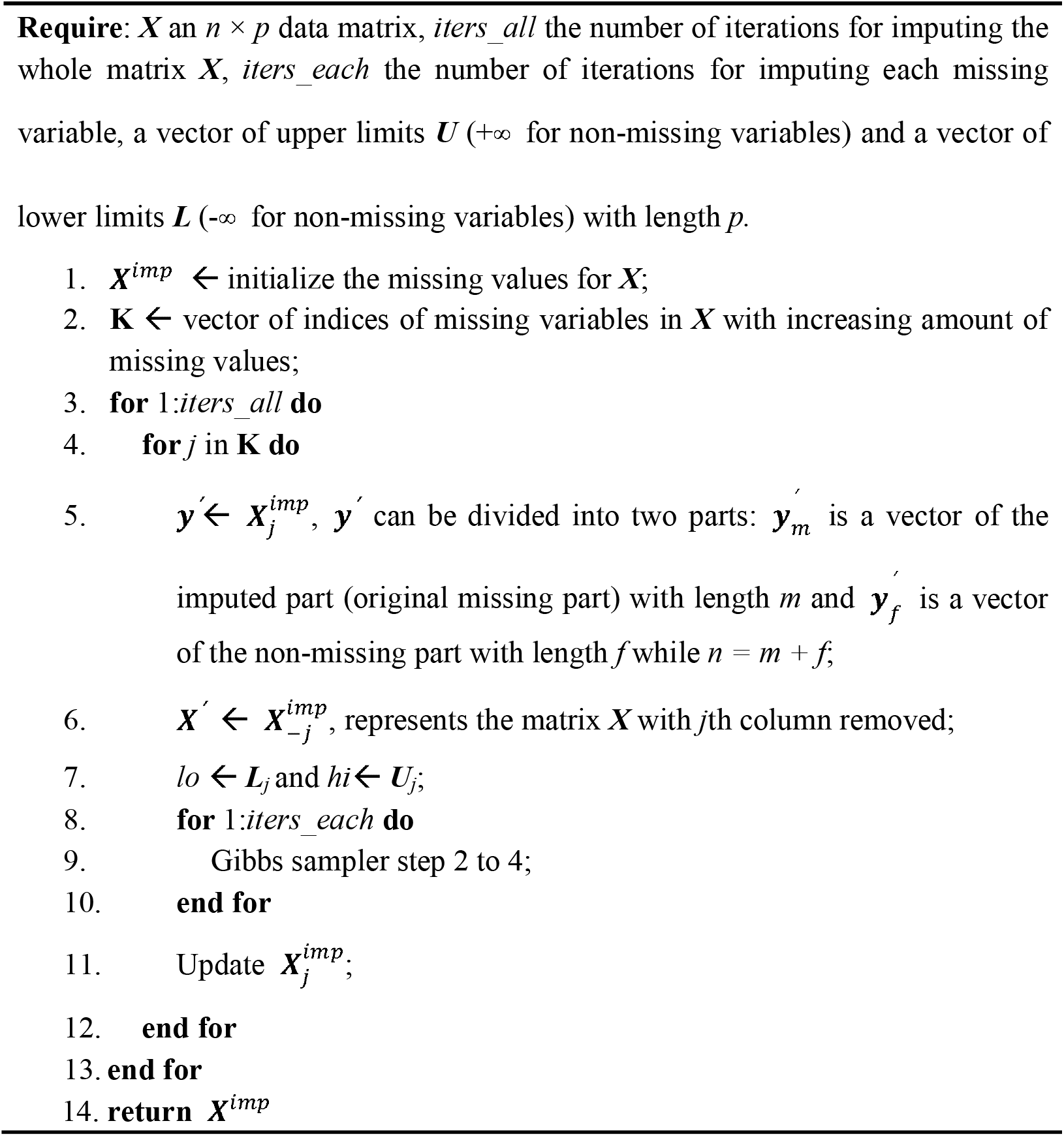

### Other imputation approaches

Other three left-censored missing imputation/substitution methods were conducted in our study for performance comparison:

- kNN-TN (Truncation *k*-nearest neighbors imputation) [21]: this method applied a Newton-Raphson (NR) optimization to estimate the truncated mean and standard deviation. Then, Pearson correlation was calculated based on standardized data followed by correlation-based kNN imputation.
- QRILC (Quantile Regression Imputation of Left-Censored data) [18,30]: this method imputes missing elements randomly drawing from a truncated distribution estimated by a quantile regression. R package *imputeLCMD* was applied for this imputation approach.
- HM (Half of the Minimum): This method replaces missing elements with half of the minimum of non-missing elements in the corresponding variable.

### Assessments of performance

The assessments of imputation performance were conducted using an imputation evaluation pipeline from our previous study with both unlabeled and labeled measurements [31], which is accessible through: https://github.com/WandeRum/MVI-evaluation. Unlabeled measurements include the NRMSE-based sum of ranks (SOR), principal component analysis (PCA)-Procrustes analysis while labeled measurements include correlation analysis for univariate results, partial least square (PLS)-Procurstes analysis. R package *vegan* was applied for Procrustes analysis [32] and *ropls* was applied for PLS analysis [33].

Furthermore, we evaluated the impacts of different imputation methods on the statistical sensitivity of detecting biological variances. On the simulation dataset, we calculated *p*-values from student’s *t*-tests between two groups from original as well as imputed datasets. We marked a set *S* as real differential variables at a significant level of *p*-cutoff (e.g. 0.05) from original simulation data, and a set *S’* as detected differential variables at the same significant level from imputed simulation data. Then we calculated the true positive rate 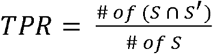 to evaluate the effects of different imputation methods in terms of detecting differential variables.

## Conclusion

A practical left-censored missing value imputation method is needed in the field of metabolomics. We develop a new imputation approach GSimp that outperforms traditional determined value substitution method (HM) and other approaches (QRILC, and kNN-TN) for MNAR situations. GSimp utilized predictive information of variables and held a truncated normal distribution for each missing element simultaneously via embedding a prediction model into the Gibbs sampler framework. With proper modifications on the parameter settings, e.g. truncation points, GSimp may be applicable to handle different types of missing values and in different-omics studies, thus deserved to be further explored in the future.

---

**S1 Fig.**
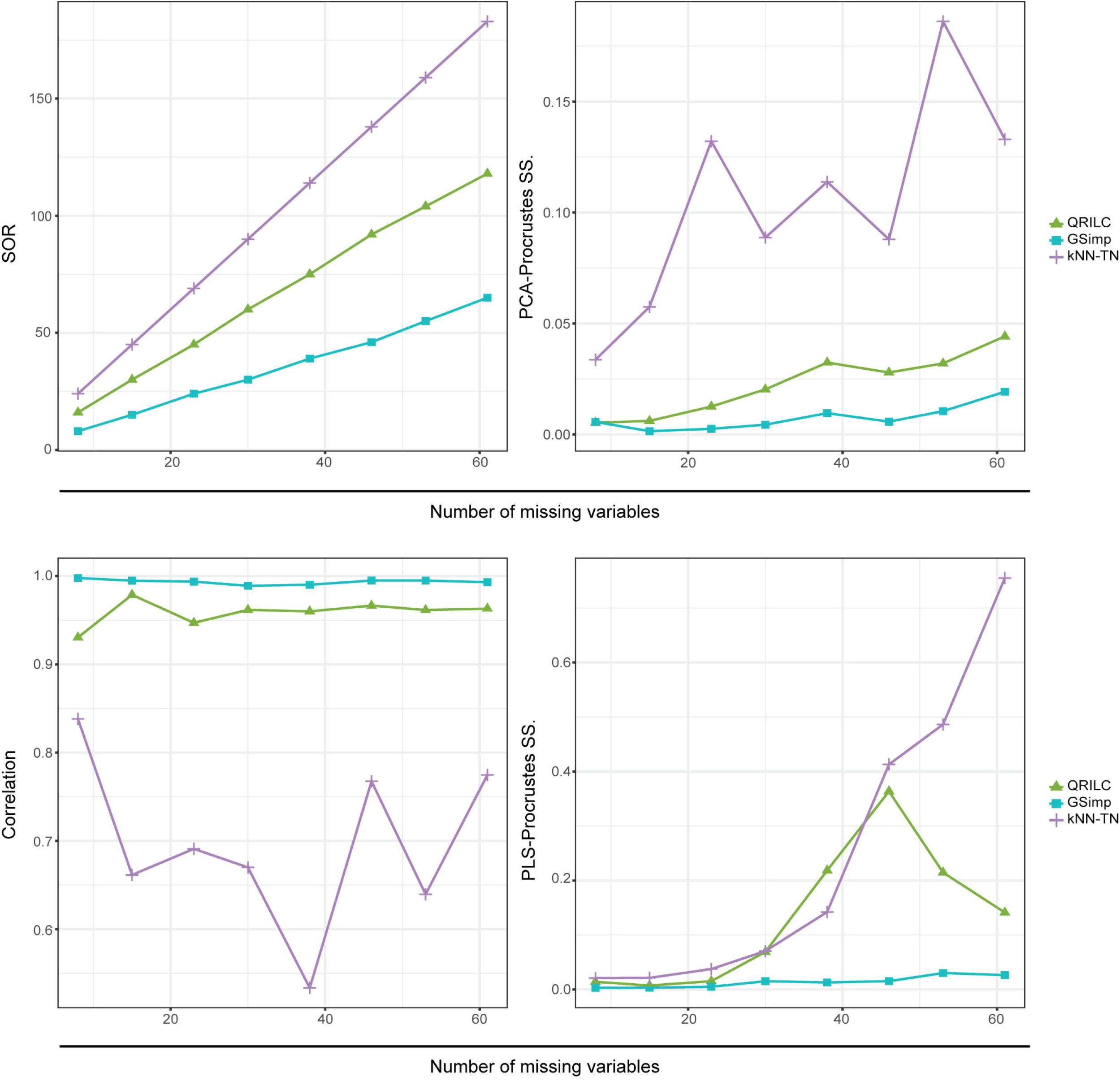
Evaluations of different imputation methods on simulation dataset. SOR (upper left), PCA-Procrustes sum of squared errors (upper right), Pearson’s correlation between log-transformed *p*-values of student’s t-tests (lower left), and PLS-Procrustes sum of squared errors (lower right) on simulation dataset along with different numbers of missing variables based on three imputation methods: QRILC (green triangle), GSimp (blue square), and kNN-TN (purple cross).

**S2 Fig.**
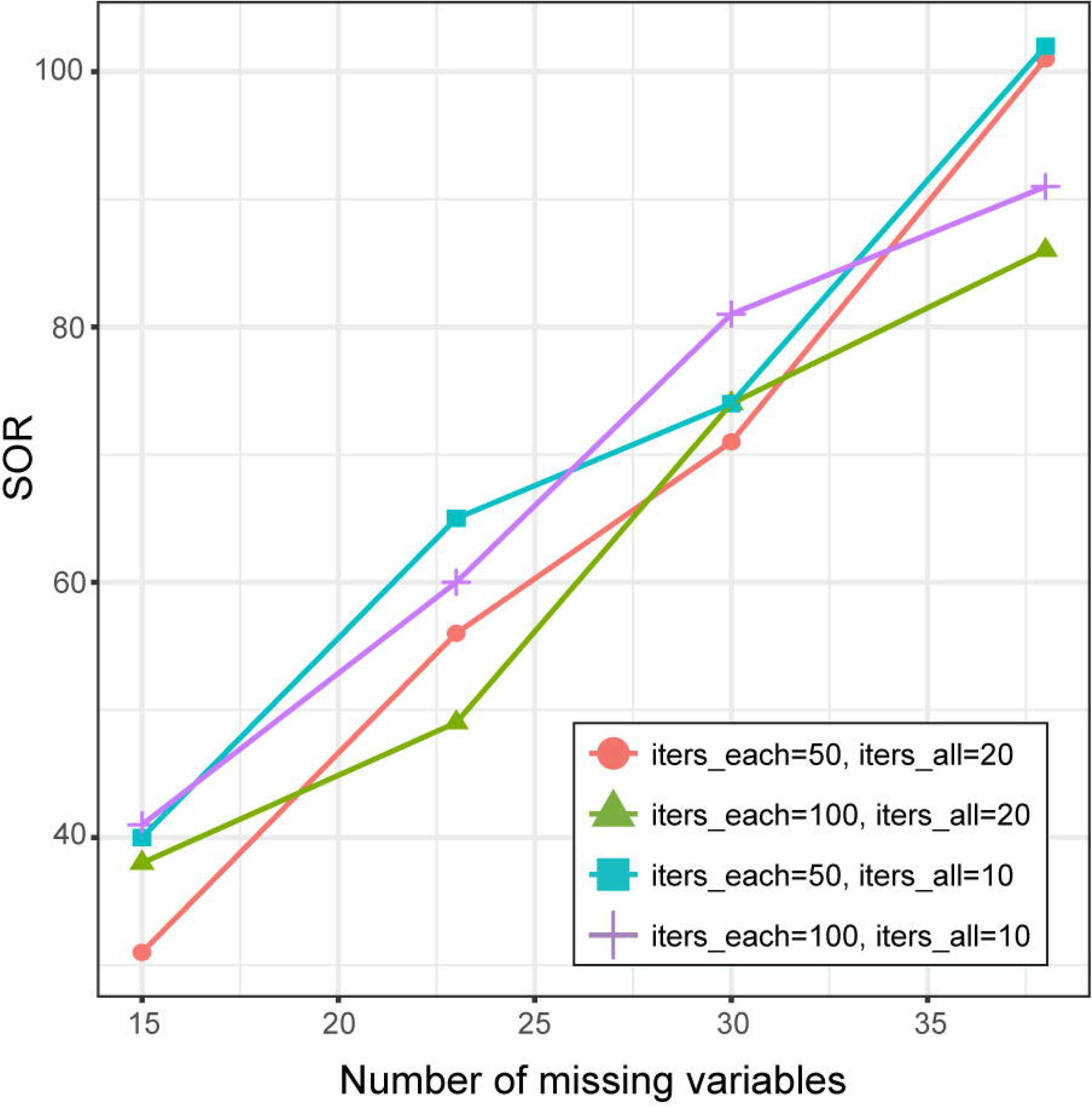
Evaluations of different numbers of iterations using GSimp on simulation dataset. SOR on simulation dataset along with different numbers of missing variables based on four different numbers of iterations: *iters_each*=50 and *iters_all*=20 (red circle), *iters_each*=100 and *iters_all*=20 (green triangle), *iters_each*=50 and *iters_all*=10 (blue square), *iters_each*=100 and *iters_all*=10 (purple cross).

## References

1. Gelman A, Hill J. Data analysis using regression and multilevel/hierarchical models [Internet]. Cambridge University Press. 2006. doi:10.2277/0521867061

2. Little RJ a, Rubin DB. Statistical Analysis with Missing Data. Statistical analysis with missing data Second edition. 2002. doi:10.2307/1533221

3. Hrydziuszko O, Viant MR. Missing values in mass spectrometry based metabolomics: An undervalued step in the data processing pipeline. Metabolomics. 2012;8: 161–174. doi:10.1007/s11306-011-0366-4

4. Guo L, Milburn M V, Ryals JA, Lonergan SC, Mitchell MW, Wulff JE, et al Plasma metabolomic profiles enhance precision medicine for volunteers of normal health. Proc Natl Acad Sci. 2015;112: E4901–E4910. doi:10.1073/pnas.1508425112

5. Liu J-J, Ghosh S, Kovalik J-P, Ching J, Choi HW, Tavintharan S, et al Profiling of plasma metabolites suggests altered mitochondrial fuel usage and remodelling of sphingolipid metabolism in individuals with type 2 diabetes and kidney disease. Kidney Int Reports. 2016;2: 470–480. doi:10.1016/j.ekir.2016.12.003

6. Butte NF, Liu Y, Zakeri IF, Mohney RP, Mehta N, Voruganti VS, et al Global metabolomic profiling targeting childhood obesity in the Hispanic population. Am J Clin Nutr. 2015;102: 256–267. doi:10.3945/ajcn.115.111872

7. Troyanskaya O, Cantor M, Sherlock G, Brown P, Hastie T, Tibshirani R, et al Missing value estimation methods for DNA microarrays. Bioinformatics. 2001;17: 520–525. doi:10.1093/bioinformatics/17.6.520

8. Hastie T, Tibshirani R, Sherlock G. Imputing missing data for gene expression arrays. Tech Report, Div Biostat Stanford Univ. 1999; 1–9.

9. Stacklies W, Redestig H, Scholz M, Walther D, Selbig J. pcaMethods - A bioconductor package providing PCA methods for incomplete data. Bioinformatics. 2007;23: 1164–1167. doi:10.1093/bioinformatics/btm069

10. Stekhoven DJ, Bühlmann P. Missforest-Non-parametric missing value imputation for mixed-type data. Bioinformatics. 2012;28: 112–118. doi:10.1093/bioinformatics/btr597

11. Mak TD, Laiakis EC, Goudarzi M, Fornace AJ. MetaboLyzer: A novel statistical workflow for analyzing postprocessed LC-MS metabolomics data. Anal Chem. 2014;86: 506–513. doi:10.1021/ac402477z

12. Katajamaa M, Miettinen J, Oresic M. MZmine: toolbox for processing and visualization of mass spectrometry based molecular profile data. Bioinformatics. 2006;22: 634–6. doi:10.1093/bioinformatics/btk039

13. Kessler N, Neuweger H, Bonte A, Langenkämper G, Niehaus K, Nattkemper TW, et al MeltDB 2.0-advances of the metabolomics software system. Bioinformatics. 2013;29: 2452–2459. doi:10.1093/bioinformatics/btt414

14. Luedemann A, Von Malotky L, Erban A, Kopka J. TagFinder: Preprocessing software for the fingerprinting and the profiling of gas chromatography-mass spectrometry based metabolome analyses. Methods Mol Biol. 2012;860: 255–286. doi:10.1007/978-1-61779-594-7_16

15. Xia J, Sinelnikov I V., Han B, Wishart DS. MetaboAnalyst 3.0-making metabolomics more meaningful. Nucleic Acids Res. 2015;43: W251–W257. doi:10.1093/nar/gkv380

16. Xia J, Psychogios N, Young N, Wishart DS. MetaboAnalyst: A web server for metabolomic data analysis and interpretation. Nucleic Acids Res. 2009;37. doi:10.1093/nar/gkp356

17. Xia J, Mandal R, Sinelnikov I V., Broadhurst D, Wishart DS. MetaboAnalyst 2.0-a comprehensive server for metabolomic data analysis. Nucleic Acids Res. 2012;40. doi:10.1093/nar/gks374

18. Lazar C, Gatto L, Ferro M, Bruley C, Burger T. Accounting for the Multiple Natures of Missing Values in Label-Free Quantitative Proteomics Data Sets to Compare Imputation Strategies. J Proteome Res. 2016;15: 1116–1125. doi:10.1021/acs.jproteome.5b00981

19. Shah JS, Rai SN, DeFilippis AP, Hill BG, Bhatnagar A, Brock GN. Distribution based nearest neighbor imputation for truncated high dimensional data with applications to pre-clinical and clinical metabolomics studies. BMC Bioinformatics. 2017;18: 114. doi:10.1186/s12859-017-1547-6

20. Lazar C, Gatto L, Ferro M, Bruley C, Burger T. Accounting for the Multiple Natures of Missing Values in Label-Free Quantitative Proteomics Data Sets to Compare Imputation Strategies. J Proteome Res. 2016;15: 1116–1125. doi:10.1021/acs.jproteome.5b00981

21. Shah JS, Rai SN, DeFilippis AP, Hill BG, Bhatnagar A, Brock GN. Distribution based nearest neighbor imputation for truncated high dimensional data with applications to pre-clinical and clinical metabolomics studies. BMC Bioinformatics. 2017;18: 114. doi:10.1186/s12859-017-1547-6

22. Ni Y, Zhao L, Yu H, Ma X, Bao Y, Rajani C, et al Circulating Unsaturated Fatty Acids Delineate the Metabolic Status of Obese Individuals. EBioMedicine. 2015;2: 1513–1522. doi:10.1016/j.ebiom.2015.09.004

23. Lei S, Huang F, Zhao A, Chen T, Chen W, Xie G, et al The ratio of dihomo-γ-linolenic acid to deoxycholic acid species is a potential biomarker for the metabolic abnormalities in obesity. FASEB J. Federation of American Societies for Experimental Biology; 2017; fj.201700055R. doi:10.1096/fj.201700055R

24. Zou H, Hastie T. Regularization and variable selection via the elastic net. J R Stat Soc Ser B Stat Methodol. 2005;67: 301–320. doi:10.1111/j.1467-9868.2005.00503.x

25. Breiman L, Friedman JH, Olshen RA, Stone CJ. Classification and Regression Trees. The Wadsworth statisticsprobability series. 1984. doi:10.1371/journal.pone.0015807

26. Breiman L. Random forests. Mach Learn. 2001;45: 5–32. doi:10.1023/A:1010933404324

27. Tibshirani R. Regression Selection and Shrinkage via the Lasso [Internet]. Journal of the Royal Statistical Society B. 1996. pp. 267–288. doi:10.2307/2346178

28. Hoerl AE, Kennard RW. Ridge Regression: Biased Estimation for Nonorthogonal Problems. Technometrics. 1970;12: 55–67. doi:10.1080/00401706.1970.10488634

29. Friedman AJ, Hastie T, Simon N, Tibshirani R, Hastie MT. Lasso and Elastic-Net Regularized Generalized Linear Models [Internet]. 2015. Available: http://www.jstatsoft.org/v33/i01/.

30. Lazar C. Imputation of left-censored missing data using QRILC method [Internet]. 2015.

31. Wei R, Wang J, Su M, Jia E, Chen T, Ni Y. Missing Value Imputation Approach for Mass Spectrometry-based Metabolomics Data. bioRxiv. 2017; Available: http://biorxiv.org/content/early/2017/08/17/171967.abstract

32. Oksanen J. Multivariate Analysis of Ecological Communities in R: vegan tutorial [Internet]. 2015.

33. Thévenot EA, Roux A, Xu Y, Ezan E, Junot C. Analysis of the Human Adult Urinary Metabolome Variations with Age, Body Mass Index, and Gender by Implementing a Comprehensive Workflow for Univariate and OPLS Statistical Analyses. J Proteome Res. 2015;14: 3322–3335. doi:10.1021/acs.jproteome.5b00354

